# A new clinical tool to predict outcome in early-stage melanoma patients

**DOI:** 10.1101/632455

**Authors:** Filippo Mancuso, Sergio Lage, Javier Rasero, José Luis Díaz-Ramón, Aintzane Apraiz, Gorka Pérez-Yarza, Pilar A. Ezkurra, Cristina Penas, Ana Sánchez-Diez, María Dolores García-Vazquez, Jesús Gardeazabal, Rosa Izu, Karmele Mujika, Jesús Cortés, Aintzane Asumendi, María Dolores Boyano

## Abstract

Around 25% of early-stage melanoma patients eventually develop metastasis. Thus, we set out to define serological biomarkers that could be used along with clinical and histopathological features of the disease to predict these events. We previously demonstrated that in stage II melanoma patients, serum levels of dermcidin (DCD) were associated with metastatic progression. Based on the relevance of the immune response on the cancer progression and the recent association of DCD with local and systemic immune response against cancer cells, serum DCD was analyzed in a new cohort of patients along with IL-4, IL-6, IL-10, IL-17A, IFNγ TGFβ and GM-CSF. We included 448 melanoma patients, 323 of whom were diagnosed as stages I-II according to AJCC. Levels of selected cytokines were determined by ELISA and Luminex and obtained data were analyzed employing Machine Learning and Kaplan-Meier techniques to define an algorithm capable of accurately classifying early-stage melanoma patients with a high and low risk of developing metastasis. The results show that in early-stage melanoma patients, serum levels of the cytokines IL-4, GM-CSF and DCD together with the Breslow thickness are those that best predict melanoma metastasis. Moreover, resulting algorithm represents a new tool to discriminate subjects with good prognosis from those with high risk for a future metastasis.

**Novelty and Impact:** We have developed a prognostic equation that considers the serum IL-4, GM-CSF and DCD levels, along with the Breslow thickness to accurately classify melanoma outcome in patients. In this sense, a rigorous follow-up is recommended for early-stage melanoma patients with a high Breslow thickness, high serum IL-4 levels and low GM-CSF and DCD levels at the time of diagnosis, given the elevated risk for these patients to develop metastasis during follow-up.

## Introduction

Early and accurate classification of patients is the cornerstone of precision medicine, intimately linked to the optimal management of cancer. This is especially relevant for melanoma, the most deadly type of skin cancer due to its high metastatic capacity and the limited therapeutic tools available to combat the spread of the disease. In fact, distant metastases are associated with median survival rates ranging from 6 to 15 months^1^. Despite all the efforts to devise prevention and detection strategies, the incidence of melanoma is expected to increase in the forthcoming years^2^, further supporting the benefits to be gained by investing in the development of predictive tools.

The prognosis of melanoma is currently assigned almost entirely on the basis on a limited set of histopathological markers^3,4^. In this context, tumor thickness is the most important histopathological characteristic included in the AJCC staging system and it is officially considered as a prognostic factor for melanoma progression in clinical practice^5,6^. However, due to the clinical and biological heterogeneity of primary melanoma, survival can vary widely even among individuals considered to be within the same stage^7,8^, highlighting the need for new prognostic tools to improve the management of primary melanoma patients^9^. Precision medicine focuses on classifying early stage melanoma patients on the basis of genetic and other biochemical features in order to identify profiles that are most likely to develop into more advanced disease stages, and to define more effective treatments for the metastatic disease^10^.

Serum is a highly accessible and valuable source of biomarkers, containing tumor and host-related factors that are correlated with tumor behavior and patient prognosis^11^. Cytokines are key mediators of the immune system with either pro-inflammatory or anti-inflammatory activity, and they are serum factors with potential value as biomarkers. In fact, cytokine profiling is providing valuable data regarding patient classification in a wide range of diseases, including cancer^12,13,14^. In terms of tumor activity, elevated Th2 cytokines (IL-4 -Interleukin-4, IL-5, IL-10 and IL-13) and decreased Th1 cytokines (IL-2, TGFβ and IFNγ − Transforming growth factor β and Interferon-γ) suppress effective spontaneous anti-tumor immunity^15,16,17^. In addition, the IL-17A pro-inflammatory cytokine has been associated with poor prognosis in some tumors^18^. GM-CSF (Granulocyte-Macrophage Colony-Stimulating Factor) is a hematopoietic growth factor that fulfills a fundamental role in macrophage and granulocyte differentiation. While classically linked to anti-tumor activities^19^ there is growing evidence that GM-CSF can also promote tumor progression^20,21,22^, supporting its inclusion in biomarker studies.

In a previous study carried out on a large group of melanoma patients, and based on serum proteomic analysis and immunoassays, we established prognostic value of serum Dermcidin (DCD) for stage II melanoma patients^23^. DCD is considered to play an important role in the cutaneous microenvironment due to its antimicrobial activity^24^. Nevertheless, DCD is not just an antimicrobial peptide as it can stimulate keratinocytes to produce cytokines though G-protein and mitogen-activated protein kinase activation^25^. These data suggest a possible relationship between *in situ* and systemic immune responses. Accordingly, the purpose of this study was to determine whether selected cytokine and DCD levels in serum could be used to develop a tool with clinical applications to improve the prognostic prediction of patients diagnosed at early stages of melanoma. To achieve this, we adopted a machine learning approach that incorporated the serum measurements of GM-CSF, IFN-γ, TGFβ1, IL-4, IL-6, IL-10, IL-17A and DCD, in conjunction with clinical-pathological features of such melanoma patients to determine the prognostic value of these parameters.

## Results

### Patient characteristics

448 melanoma patients were included in this study (187 male, 261 female), with a median age at diagnosis of 56 years (95% CI 54.0-60.0: Table 1). Melanoma was most often diagnosed in patients’ trunks (158 patients), followed by the lower limb (121 patients). Head and neck, upper limb and acral locations for melanomas were identified in 69, 49 and 34 patients, respectively. A large proportion of the lesions had a Breslow thickness less than 1 mm (231 melanomas) and between 1.01-4.00 mm (131 melanomas). Staging was based on the AJCC system^7^ and most patients were diagnosed as stage I or II (224 and 99 patients, respectively), while only 38 were considered to be at a stage related with metastasis, stage III (30) or IV (8). However, 119 (27%) of the 448 patients recruited developed metastasis, including those with spread disease at the moment of diagnosis and those who suffered from disease recurrence during the follow-up. Distant and ganglion metastases were the main subtypes detected. Considering patients at AJCC stages I and II as early-stage melanoma patients (323 patients), sex, age and tumor location frequency were similar to the whole group and 84 of these early-stage melanoma patients (25.6%) developed metastasis during the first years of the follow-up. In fact, the median interval from the removal of the primary tumor until the diagnosis of a metastasis was 1.9 years. By contrast, 239 patients remained disease-free (without recurrence or metastasis) and the median follow-up of these patients was 4.5 years.

**Table 1.** Patient characteristics.

| **Characteristic** | **Number (m/f)** |
| --- | --- |
| **Melanomas** | 448 (187/261) |
| Disease-free | 329 (136/193) |
| Metastasis | 119 (51/68) |
| **Tumor location** |  |
| Head/neck | 69 (33/36) |
| Trunk | 158 (98/60) |
| Upper limb | 49 (16/33) |
| Lower limb | 121 (22/99) |
| Hand/foot | 34 (10/24) |
| Others | 7 (1/6) |
| Unknown | 10 (7/3) |
| **Stage<sup>1</sup>** |  |
| 0 (*in situ*) | 78 (29/49) |
| I | 224 (94/130) |
| II | 99 (40/59) |
| III | 30 (16/14) |
| IV | 8 (4/4) |
| Unknown | 9 (4/5) |
| **Breslow Thickness (mm)** |  |
| ≤ 1.00 | 231 (98/133) |
| 1.01-4.00 | 131 (43/88) |
| > 4.01 | 41 (25/16) |
| Unknown | 45 (21/24) |

<sup>1</sup>The American Joint Committee of Cancer (AJCC) staging system for melanomas was used. Abbreviations: m/f, males/females.

### Analysis of serum GM-CSF, IL-4, IL-6, IL-10, IL-17A, IFN-γ, TGFβ and DCD

The amount of GM-CSF, IL4, IL-6, IL-10, IL-17A and TGFβ detected in serum was independent of the age of the melanoma patients and it did not vary between the sexes. However, there were significant differences between the sexes in the levels of IFNγ and DCD (|δ|= 0.2, *p*_FDR_<0.01 and |δ|= 0.2, *p*_FDR_<0.01, respectively: data not shown). Of the proteins studied, the median serum level in the melanoma patients at the time of diagnosis was considered according to the stage of the tumor (Table 2). There appeared to be no significant differences in the median serum levels in patients of different AJCC stages, nor were any differences found between the distinct histological subtypes of melanoma (data not shown).

**Table 2.**
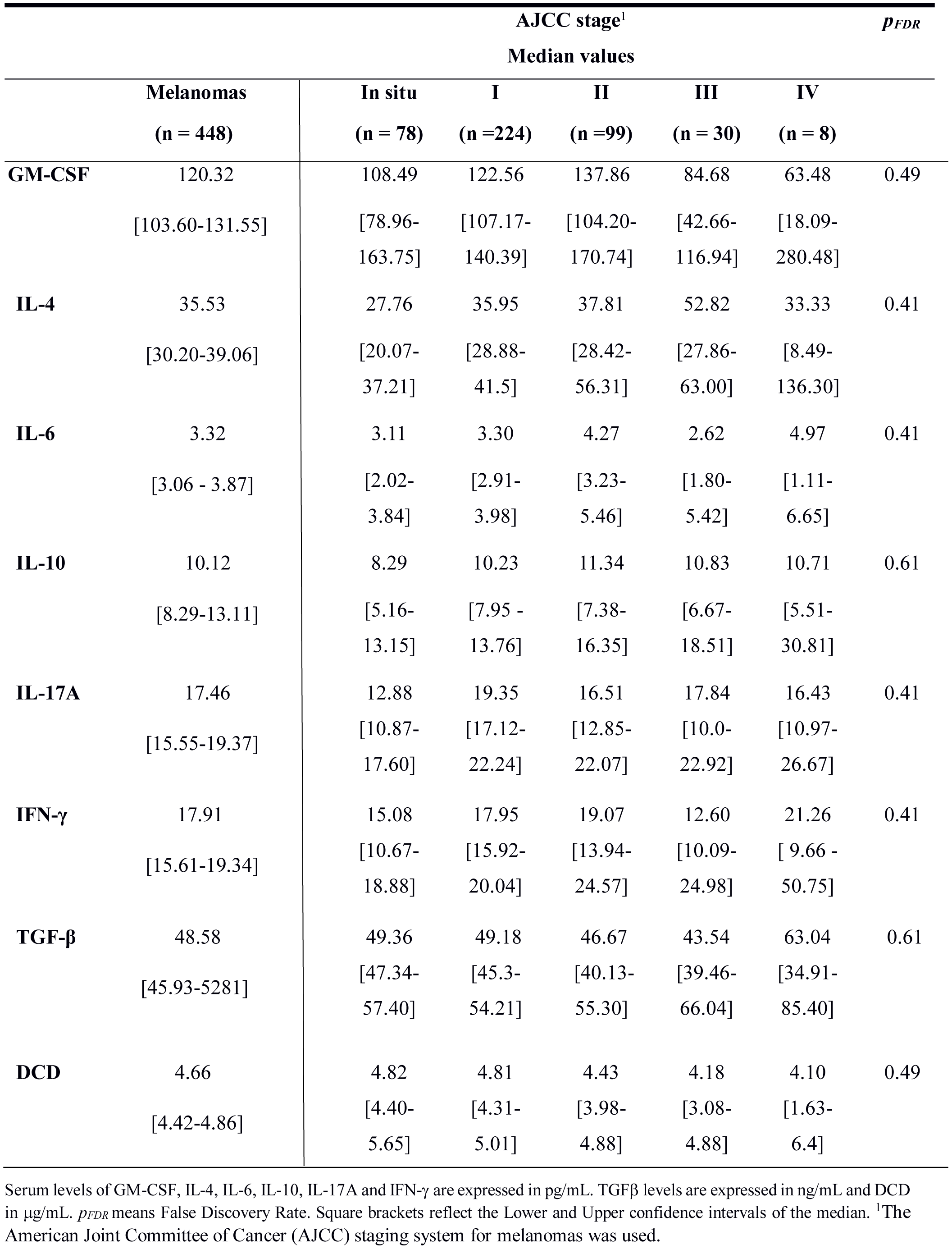
Cytokines and DCD serum levels in melanoma patients.

To analyze the prognostic value of these proteins, we assessed the serum cytokine and DCD levels in melanoma patients diagnosed at stages I/II and that remained disease-free at the end of the follow-up period, comparing these with those in patients who developed metastasis. There were 84 of 323 (25.6%) stage I/II melanoma patients who developed metastasis during the follow-up and there appeared to be a significant difference in the serum IL-4 and IL-6 levels between these two groups of patients, and associated with a moderate effect size (|δ|= 0.30 *p*_*FDR*_ <0.01 and |δ|= 0.20 *p*_*FDR*_ = 0.04: Table 3). At the time of diagnosis, the serum IL-4 levels of patients who developed metastasis doubled those observed in patients who remained disease-free (62.27 pg/mL *versus* 31.96 pg/mL, respectively, p<0.01). In addition, there was a significant difference in the serum IL-6 levels between the two groups of patients (4.71 pg/mL *versus* 3.29 pg/mL respectively, p<0.04). No significant differences in serum IL-10, IL-17A, IFN-γ, TGFβ and DCD were observed.

**Table 3.** Comparison of serum cytokine and DCD levels between AJCC stage I and II patients who were disease-free or developed metastasis during follow-up.

| Disease progression | | | (\|$\delta$\|, $p_{FDR}$) |
| --- | --- | --- | --- |
| Median Values |  |  |  |
|  | Disease-free | Metastasis |  |
| GM-CSF | 121.39 | 131.55 | (0.03, 0.72) |
|  | [103.17-138.63] | [101.01-153.24] |  |
| IL-4 | 31.96 | 62.27 | (0.30, <0.01) |
|  | [27.76-38.06] | [39.06-92.21] |  |
| IL-6 | 3.29 | 4.71 | (0.20, 0.04) |
|  | [2.82-3.87] | [3.3- 5.9] |  |
| IL-10 | 11.23 | 10.03 | (0.05, 0.67) |
|  | [7.89-14.83] | [7.78 - 15.31] |  |
| IL-17A | 17.88 | 20.09 | (0.04, 0.67) |
|  | [16.33-20.06] | [14.89-24.33] |  |
| IFN-γ | 17.23 | 22.40 | (0.12, 0.37) |
|  | [14.84-19.34] | [16.99-26.91] |  |
| TGF-β | 49.71 | 48.18 | (0.04, 0.67) |
|  | [45.32-53.68] | [40.13-60.44] |  |
| DCD | 4.78 | 4.38 | (0.07, 0.67) |
|  | [4.39-5.01] | [3.92-4.84] |  |

Serum levels of GM-CSF, IL-4, IL-6, IL-10, IL-17A and IFN-γ are expressed in pg/mL. TGFβ levels are expressed in ng/mL and DCD in $\mu$ g/mL. | $\delta$ |= Cliff's Delta and $p_{FDR}$ means False Discovery Rate. Square brackets reflect the Lower and Upper confidence intervals for the median.

### Prognostic power of the melanoma markers

The performance of the different classifiers was assessed for the subpopulation of subjects at AJCC stages I and II (Table S1). In all the three domains, a LR classifier exhibited the best performance through ROC area, and the most generalizable results reflected by the smallest gap between the training and test scores. The Breslow thickness represents a biomarker of melanoma metastasis that correctly classified 73% of the patients and generating 83% of the ROC area (Figure 1A). Although the serum levels had a poorer prognostic value, when combined with the Breslow thickness they significantly improved the cross-validated performance of this biomarker, exceeding a balanced accuracy of 80% (Wilcoxon signed-rank test, p<0.01) and a ROC area close to the 90% (Wilcoxon signed-rank test, p<0.01). Furthermore, these data clearly pointed to the cytokines IL-4, GM-CSF and DCD as the most powerful biomarkers in predicting melanoma metastasis in conjunction with the Breslow thickness. Indeed, these parameters were selected at least the 80% of the times in all the partitions after the feature selection process (see panel B of Figure 1). This subset of variables was followed by the IL-10 being 66% of the times selected and the IL-6, IL-17A, IFN-γ and TGFβ below the 50%.

**Figure 1.**
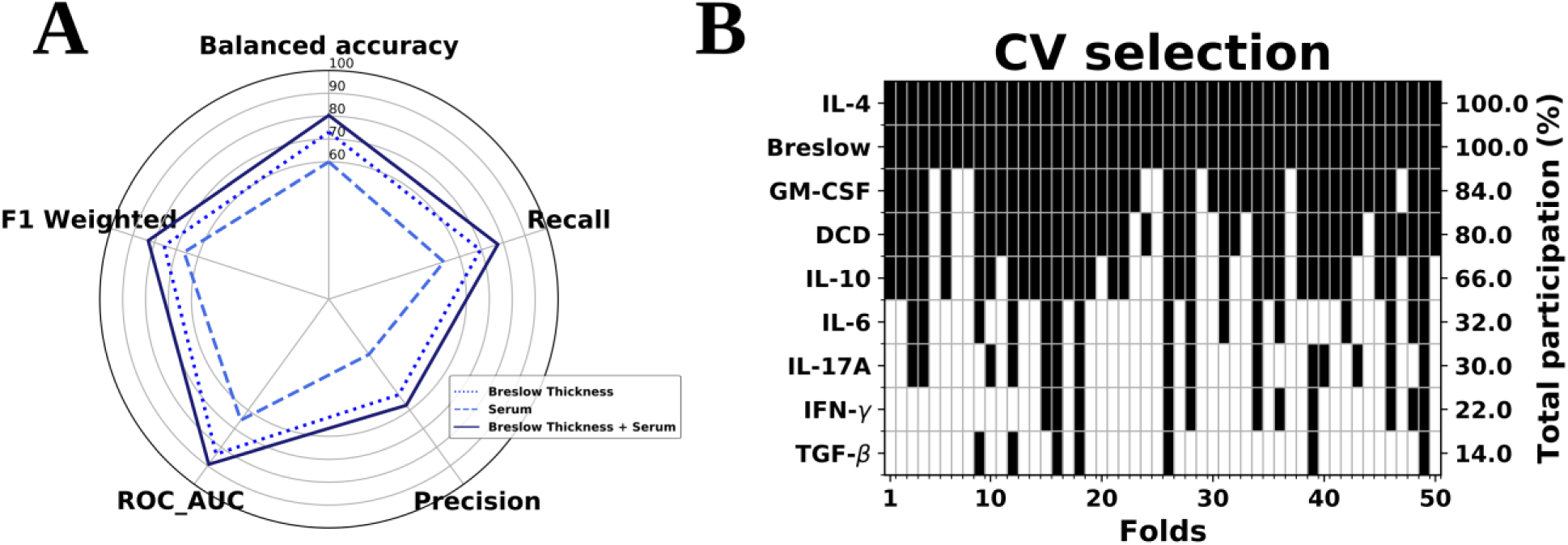
Logistic Regression analysis. A) Classification of the three variable domains considered: Breslow thickness, Cytokines and DCD serum variables. B) In the scenario combining histological and serum variables, their participation across the folds provided by the feature selection step in the inner cross-validation loop. Black colors in each column denote the predictors that were included in the final logistic regression model in each of these folds. Data from the early-stage melanoma cohort (n=323).

Notably, the decile distribution for these potential biomarkers exhibited a clear tendency to separate between subjects in stages I and II who developed metastasis and those who remained disease-free (panel A in Figure 2). Both the distributions of the serum IL-4 levels and the Breslow thickness were higher in the metastatic subpopulation, whereas this tendency switched towards lower levels of GM-CSF. Moreover, when the differences in the distribution of these variables was addressed by means of the shift function, their predictive power was clearly evident, especially that of the Breslow thickness where the separation of the melanoma outcome was significant across its whole spectrum. This was followed by that of IL-4, which began to display a significant separation around the median, whereas GM-CSF started to discriminate these subpopulations above its 8^th^ decile (panel B in Figure 2). For DCD, nevertheless, this class separating tendency is not as evident (see Figure S2), which may denote a synergetic role emerging beyond the univariate scenario. This also seems to be the same for the rest of variables of interest (Figure S2).

**Figure 2.**
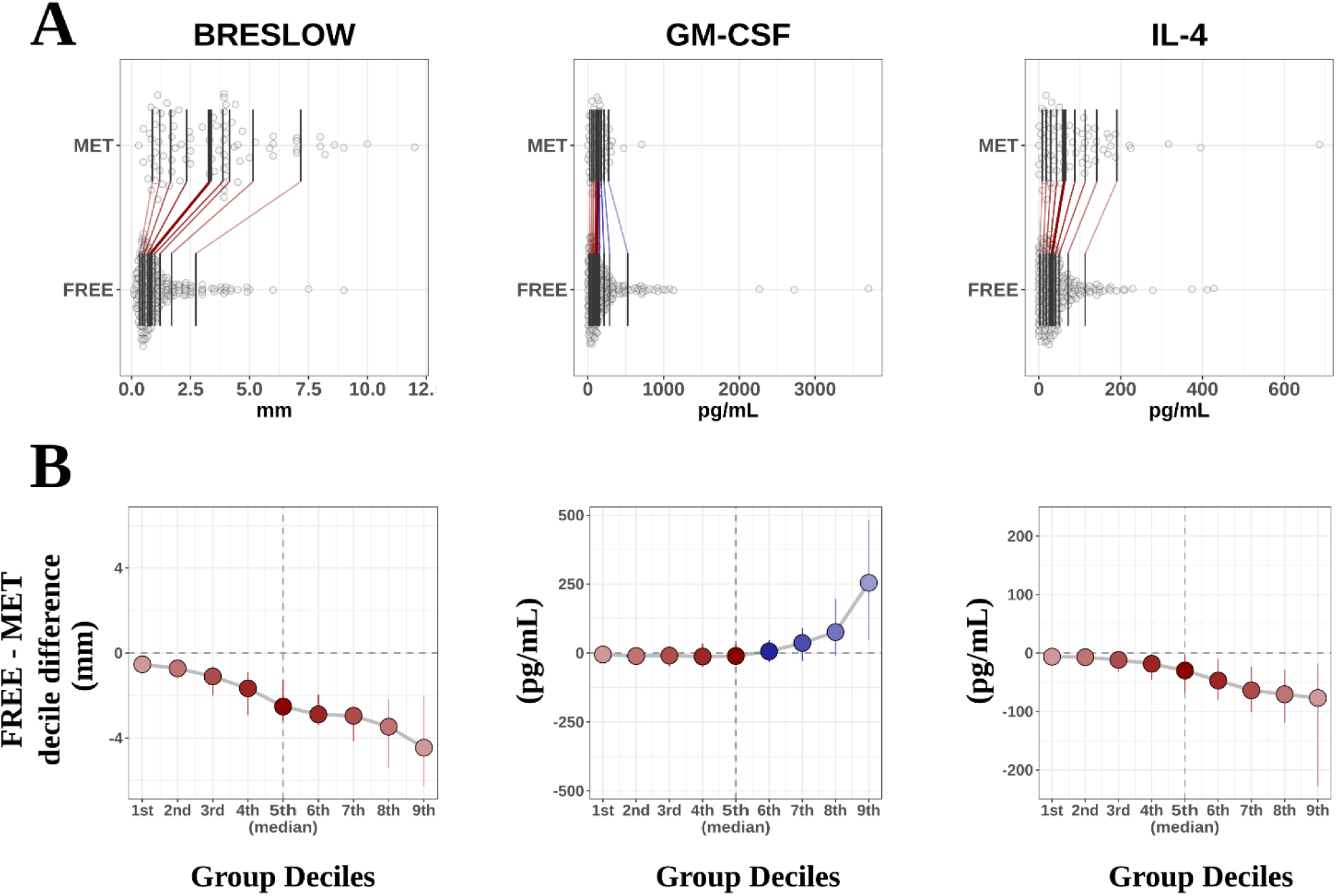
A) Decile distribution of metastatic and disease-free subjects for the Breslow thickness, GM-CSF and IL-4. B) For this subset of features, the shift function displaying the difference between the deciles in both subgroups of subjects. Positive values of the shift function are in blue, corresponding to larger decile values in the disease-free group than in the metastatic group, while red values illustrate the opposite scenario.

More importantly, these findings can be easily incorporated into clinical protocols by providing a general optimum cut-off from the data from which a prediction of metastasis can be performed. Using the same subpopulation of subjects (I/II melanoma patients), we fitted the entire data using the best classifier and the subset of features found previously (i.e.: a LR classifier with the features Breslow thickness, GM-CSF, IL-4 and DCD), and we computed the optimal point on the ROC curve that corresponded in this case to FPR (1-Specificity) = 0.11 and TPR (Sensitivity) = 0.79 (see panel A, Figure 3). This point defines a critical threshold that allows us to separate subjects in terms of their prognosis, which for our classifier can be easily translated into a constraint as follows:

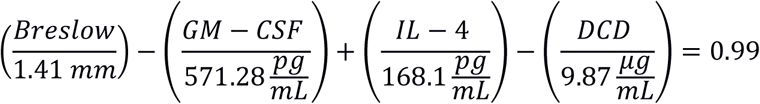

**Figure 3.**
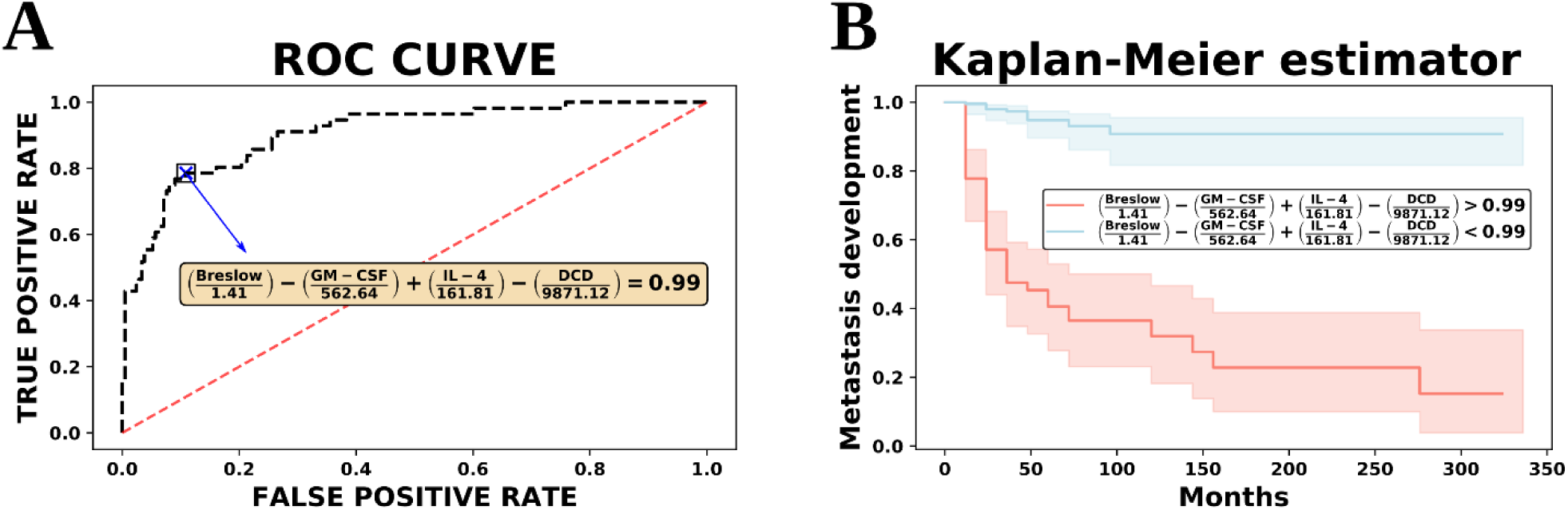
A) ROC curve from the whole stage I/II dataset. The optimal cut-off point on this curve defines a plane that maximally separates metastatic and disease-free progression. The best subset of biomarkers corresponds to Breslow thickness, IL-4, GM-CSF and DCD. B) Kaplan-Meier analysis. The cut-off plane provides a condition to significantly separate subjects with a worse prognosis from those with a better prognosis.

This equation therefore defines a hyperplane in our feature space such that any subject lying above it is classified as metastatic and those below it is considered disease-free. Furthermore, subjects stratified with respect to this critical threshold could be differentiated by their probability of eventually developing metastasis during the follow-up period (Kaplan-Meier log rank test *p* < 0.001, as shown in panel B of Figure 3).

Remarkably, we found a *prognostic plane* involving the serum levels of IL-4, GM-CSF and DCD in conjunction with the Breslow thickness that could accurately classify subjects according to their melanoma outcome. This equation could be easily translated to a clinical setting and inspecting the signs of the coefficients in this equation, we can clearly see that an increase in IL-4 and the Breslow thickness tend to shift subjects above this plane, indicating a worse prognosis, whereas GM-CSF and DCD levels act in the opposite direction.

Finally, in order to account for a possible confounding effect in the present findings, we repeated the previous analyses considering also the age and sex (after one-hot-encoding into male and female categories) as possible covariates in the full feature matrix used to train the Logistic Regression. Their inclusion increases the balanced accuracy rate to 81.60% and the precision to 61.57%, but both variables are still behind in importance in comparison with the Breslow thickness and the most powerful serum variables (see Figure S3). Furthermore, if one attempts to incorporate age and sex to the rule provided by the prognostic equation given above, the improvement in fitting is overcome by the addition of complexity in the model, which leads to an increase in the Bayes Information Criterion (BIC) from 337.17 to 355.73, providing a very strong evidence against their inclusion (Bayes Factor > 150)^26^.

## Discussion

An accurate diagnosis is an essential first step in cancer management. Most melanoma cases are detected at early disease stages and when possible, excision biopsy is the selected procedure to treat suspicious melanocytic lesions. According to the AJCC classification, Breslow thickness, together with the ulceration and mitotic index, are important variables that should be considered in tumor staging (https://cancerstaging.org/). In addition, several histological biomarkers (e.g. Melan-A, Pmel) are routinely employed for diagnostic porpoises^27^. In this regard, important efforts are being made in order to achieve less invasive techniques or to improve the accuracy of diagnostic markers^28^. Indeed, early and precise prognostic markers are urgently needed for melanoma due to its strong metastatic capacity, particularly given the low survival rate of metastatic patients and the 5-year relapse free rate of 56%, even when melanoma is detected at early stages (stage II)^1,28^. Highest recurrence rate is observed within 2-3 years after surgical treatment while recurrence probability decreases to less than 5% in patients with treated stage I-III melanomas and 5 years of disease-free follow-up^29^. According to these figures, intense medical monitoring should be implemented in the first 2-3 years after treatment, even for stage II cases, representing an important medical and economic burden. Therefore, it would be useful if early patients could be rapidly classified into high or low recurrence risk groups when contemplating efficient and sustainable personalized follow-up programs. Moreover, to assuage the unpredictable clinical behavior of melanoma much research has focused on the discovery of prognostic factors to improve the prognostic accuracy for this type of skin cancer^3^. In this regard, our study focused on the discovery of prognostic biomarkers capable of evaluating the metastatic risk of patients identified at early stages of the disease (stage I-II). Moreover, we defined a clinically applicable mathematical tool to accurately classify such melanoma patients.

Currently, predicting patient outcome mainly relies on staging based on the histopathological parameters described previously, while treatment options are often based on the BRAF mutation^30^. Nevertheless, patient monitoring, especially upon surgical removal of the primary tumor, requires other variables to be analyzed. As a systemic system for information transfer, serum represents a complex but accessible sensor. To date, LDH has been one and perhaps the only clinical serological biomarker for melanoma, with increasing values interpreted as disease progression. However, an increase in serum LDH levels may also occur in other settings, which means employing some caution before reaching any conclusions^30^.

Our previous attempt to identify novel serological prognostic markers identified a threshold for serological DCD that was associated with a poor prognosis value for melanoma patients diagnosed specifically at AJCC stage II^23^. Consistent with this finding, DCD, a major human antimicrobial peptide in human skin^24,25^ was also recently proposed as a serological marker for the diagnosis and staging of hepatocellular carcinoma^31^.

On the other hand, it has been described that DCD is expressed in melanoma cell lines, and it could be involved in autophagy and apoptotic cell death. Nowadays, the underlying molecular mechanisms and the true clinical-pathological relevance of this is not fully understood^32^.

The current study including new melanoma patients revealed that DCD is a marker of metastatic progression, although other serological parameters appear to have greater predictive potential than DCD, such as IL-4 and GM-CSF. These differences with our previous study^23^ may be due to the patient stratification, as both stage I and stage II melanoma patients were included in separate groups.

Serological cytokines reflect the general immunological state of the body, offering information regarding the cytokines released by tumors and those that accumulate in the tumor microenvironment^33^. The melanoma microenvironment contains stromal cells and immune cells like T or B lymphocytes, NK cells or tumor-associated macrophages^33,34^. Most of these cells secrete cytokines that may play a key role in inhibiting or promoting tumor progression^35^. The pro-inflammatory cytokines IL-4 and IL-6, produced either by host immune cells or tumor cells themselves, are associated with tumor malignancy in patients and animal cancer models^36,37,38^. At the cutaneous level, keratinocytes secrete IL-6 in order to enhance T cell-mediated antitumor activity and therefore, high IL-6 levels are considered a marker for immune system upregulation^37,38^. IL-4 is the most important Th2 cytokine, and it is mainly produced by activated T cells, mast cells, basophils and eosinophils in order to regulate lymphocyte proliferation and survival^37^. Interestingly, elevated serum IL-6 was correlated with a poor prognosis in melanoma, while IL-4 is thought to promote the proliferation and survival of several cancer cells^39,40,41^. In line with previous findings, here early stage (I or II) melanoma patients that developed metastasis had significantly higher levels of serum IL-4 and IL-6 than patients who did not develop metastasis during the follow-up.

The Breslow thickness is a crucial prognostic factor, with substantial evidence confirming a direct relationship between Breslow thickness and survival^6^. Accordingly, we show that Breslow thickness is an important risk factor for the malignant progression of melanoma. Nonetheless, a significant increase in the predictive power of Breslow thickness was achieved by combining it with data regarding serum IL-4, GM-CSF and DCD, resulting in the development of an algorithm to identify early stage melanoma patients with a high risk of developing metastasis during the follow-up. According to this algorithm, a high Breslow thickness and serum IL-4 levels in early stage melanoma patients are associated with a poor prognosis, whereas GM-CSF and DCD levels decrease in patients in whom the disease outcome is poor. These results are consistent with other studies describing an antitumor effect of GM-CSF and DCD^42, 23^. Our data also revealed the importance of IL-10 and IL-6 in predicting metastatic progression, although they did not appear to add additional information to the predictive equation involving the Breslow thickness, and serum levels of IL-4, GM-CSF and DCD.

In summary, the use of machine learning techniques has helped to define an algorithm capable of accurately classifying early stage melanoma patients with a high or low risk of developing metastasis. The equation generated took into account the serum IL-4, GM-CSF and DCD levels, and the Breslow index, and it could stratify melanoma patients to be triaged at the time of diagnosis and initial surgery, or it could also be used clinically to determine whether stage in I or II melanoma patients should receive adjuvant therapy to prevent metastatic progression.

## Methods

### Patients

A total of 448 patients were recruited at the Dermatology Units at the Basurto and Cruces University Hospitals between 2008 and 2016, of whom 261 (58%) were women and 187 (42%) men. Inclusion criteria were: 1) a histologically confirmed diagnosis of malignant melanoma; 2) no treatment except primary surgery; 3) no infection as judged by clinical evaluation and the absence of increased infectious parameters in the blood.

After surgery of the primary tumor, clinical check-ups were scheduled every three months for the first two years of the follow-up, and every six months thereafter, until a five-year follow-up had been completed. Annual revisions were then scheduled up to the tenth year post-surgery. The patients who developed metastasis during the follow-up period were again examined every three months for two years after metastasis had been diagnosed. The presence or absence of metastasis was assessed in all patients by physical examination, as well as through laboratory and radiological testing (X-rays and/or computed tomography –CT-scanning). Metastases were detected in 119 of the 448 melanoma patients (27%) during this study, including those in whom the disease had spread at the moment of diagnosis, and 79 patients died due to distant metastasis (40 women and 39 men).

Disease stages were classified according to the AJCC^7^. The clinical and diagnostic data for each patient were collected retrospectively from centralized electronic and/or paper medical records. For the statistical prediction analysis, only melanoma patients at early disease stages (I and II) were included, and the inclusion of the “disease-free” group required a minimum tracking of 2 years.

The study was approved by the Euskadi Ethics Committee (reference 16-99) and written informed consents were obtained from all the subjects. The serum samples collected were stored at −80 °C at the Basque Biobank until use (https://www.biobancovasco.org/).

### Serum samples

Venous blood samples were drawn at the time of diagnosis and these samples were used to obtain serum following the protocol established at the Basque BioBank for Research. Accordingly, blood samples were allowed to clot at room temperature for at least 30 minutes and they were then centrifuged at 1000 g for 10 minutes. The serum was removed and was it subsequently divided into 500 µL aliquots, which were stored at −80 °C until use.

### Quantification of Granulocyte-Macrophage Colony-Stimulating Factor (GM-CSF), Interferon-γ (IFN-γ), Transforming Growth Factor beta (TGF-β1) and Interleukins (IL) 4, 6, 10 and 17A in serum

Upon reception, the serum samples were divided into 25 μL aliquots to avoid multiple freeze/thaw cycles, and the GM-CSF, IFN-γ, IL-4, 6, 10, 17A, and TGF-β1 in the samples was measured using magnetic bead-based multiple immunoassays (MILLIPLEX^®^ MAP kit, Human High Sensitivity T Cell Magnetic Bead Panel: EMD Millipore Corporation, Germany). Each assay included two calibration curves for each of the proteins to be measured (calibration ranges: GM-CSF 1.22-5,000 pg/mL; IFN-γ 0.61-2,500 pg/mL; IL-4 1.83-7,500 pg/mL; IL-6 0.18-750 pg/mL; IL-10 1.46-6,000 pg/mL; IL-17A 0.73-3,000 pg/mL; and TGF-β1 9.8-10,000 ng/mL), with 8 calibration points in each curve. Two low and two high quality controls were also included in the assays. In the case of TGF-β1, serum samples were treated with 1N HCl, diluting the samples 1:4 and then adding 2 µL of 1.0 N HCl before incubating the mixture for 1 h at room temperature. The samples were then further diluted 1:6 in assay buffer to achieve a final dilution of 1:30. In the assays we followed the protocol established by the manufacturer. The plates were read on a Luminex 100™ apparatus (Luminex Corporation, The Netherlands): 50 events per bead; 150 µL of sample (or 100 µL in the case of TGFβ1); gate settings from 8,000 to 15,000; reported gain as default and time out 100 s. The serum concentration of each protein was calculated through a 5-parameter logistic curve-fitting method using the xPONENT^®^ software (Luminex Corporation).

### DCD Quantification in serum

Serum DCD was measured with an ELISA kit (Cusabio Biotech Co., Ltd, Houston, US) according to manufacturer instructions as we have previously described^23^.

The optical density was determined on a microplate reader (Synergy HT, Biotek Instruments, Inc., Vermont, USA) set to 450 and 540 nm. Readings at 540 nm were subtracted from those obtained at 450 nm to correct for optical imperfections in the plate, and the serum DCD levels were calculated using the Gen5 software (2005, Biotek Instruments, Inc., Vermont, USA) with a 4-parameter logistic curve-fitting.

### Statistical analysis

Variables of interest clearly deviated from a normal distribution as assessed both visually and by means of a Shapiro-Wilk test. As a consequence, all descriptive statistic was expressed as the median along with the 95% confidence interval (CI), computed by bootstrap resampling in which 10,000 samples were extracted with replacement for each variable from the original data and calculating the 95% percentile interval. Inter-group comparisons were carried out using the Kruskal-Wallis test when more than two groups were involved and a two-sided Mann–Whitney U test when only two groups were compared. In the latter case, in addition to the p-values, the effect sizes were reported, measured through the absolute Cliff’s Delta value^43^, which estimates the difference between the probability that a value from one of the groups is higher than that value from the other group, and vice versa. The p-values were corrected for multiple comparisons by controlling the False Discovery Rate (FDR) using the Benjamini-Hochberg procedure^44^ and those whose significance level was below the threshold of were considered significant. Likewise, further inspection of statistical significance was addressed by means of the shift function^45^ as implemented in the *rogme* R package^46^, where deciles are compared using a Harrell-Davis estimator, levels of confidence computed by a bootstrap estimation and the type I error controlled to remain around 0.05 across all the decile comparisons.

### Machine learning analysis

A machine learning analysis was performed in order to assess the power of the data to correctly classify the prognosis of melanoma patients. We focused on patients diagnosed at stage I and II due to the clinical relevance of early metastasis prediction, of which there were 323 subjects in our cohort. Among these patients 244 remained disease free and 84 developed metastasis during the follow-up period. The predictive power of different biomarkers was inspected in three different variable domains: The Histological domain, represented by the Breslow thickness; the Serum domain, which involved all the serum variables indicated above; and a Multi-modal domain, a conjunction of the variables from the two previous domains. Missing information was imputed by removing instances containing unknown components, which reduced the input data to 211 disease-free and 56 metastatic samples, respectively. Subsequently, a nested cross-validation was employed to assess both the optimization and generalization of the model. In the outer loop, a 10-fold cross-validation repeated 5 times with different randomization seeds was performed to estimate the generalization error of the model. In the inner loop, a stratified 10-fold cross-validation was implemented for model optimization, which involved the tuning of a pipeline assembled by robust scaling of the data, random over sampling of the class minority, feature selection based on importance weights and classifier hyperparameter fitting. A scheme of this workflow is shown in Figure S1 (Supplementary Material).

Classification scores were computed using a battery of five classification algorithms: Logistic Regression (LR) with a L2 regularization term; Support Vector Machine (SVM) with a radial basis kernel; a Decision Tree (DT); Gaussian Naive Bayes (NB) classifier; and the K-Nearest Neighbors vote (KNN) algorithm. All the different hyperparameters of the mentioned classifiers and the level of shrinkage and the number of features to select were tuned by an exhaustive grid search within the inner loop. Finally, for each classifier, the balanced accuracy, which calculates the raw accuracy of each sample weighted by the inverse prevalence of its true class, the Precision, Recall and F1-score were reported. In addition, a Receiver Operating Characteristic (ROC) Curve was computed, such that the area under this curve (AUC) provides a measure to evaluate the classifier quality. The classifier with the highest ROC area was finally considered the most efficient one.

All the machine learning analysis was performed using *scikit-learn*, a library for machine learning written in python^47^ and *Imbalanced-learn*, a Python toolbox to “tackle the curse of imbalanced datasets in Machine Learning”^48^.

### Survival analysis

Once the best algorithm and subset of biomarkers to reflect the evolution of metastasis had been found, we used this combination to fit the entire stage I/II sub-population, allowing us to compute the ROC curve. Subsequently, the optimal cut-off point on this curve was determined using the Index of Union method, which corresponds to computing the value where the sensitivity and specificity are the closest to the AUC, and the absolute difference between the specificity and sensitivity is minimal^49^. This cut-off point allows us to define a class partition criterion, which separates subjects with a high probability of developing metastasis from those with a low probability, as witnessed through a Kaplan-Meier estimator implemented in a *lifelines* library in Python^50^.

### Ethics approval and consent to participate

The study was conducted in accordance with the Declaration of Helsinki principles. It was approved by the Euskadi Ethics Committee (reference 16-99) and written informed consents were obtained from all the subjects. The serum samples collected were stored at −80 °C at the Basque Biobank until use (https://www.biobancovasco.org/).

## Acknowledgements

We are grateful to the Basque Biobank for providing the serum samples. We are also most grateful to Drs Arantza Arrieta and Natalia Maruri (Cruces University Hospital) for their technical support with the serum marker detection. This work was supported by grants from the Basque Government (KK2016-036 and KK2017-041 to MDB), UPV/EHU (GIU17/066 to MDB), H2020-ESCEL JTI (15/01 to MDB) and MINECO (PCIN-2015-241 to MDB).

## Data Availability

The datasets generated during and/or analysed during the current study are not publicly available to avoid their potential misuse or misinterpretation, but they are available from the corresponding author on reasonable request.

## Code Availability

Code used for the analysis and plots is available in https://github.com/jrasero/citosines-melanoma.

## Author Contributions statement

MDB, AAs carried out the project administration, conceptualization, work supervision, writing - original draft preparation and review and editing- and funding acquisition.

FM, SL and MDGV performed the serological examinations, methodology.

JR and JMC have been the major contributors in statistical analysis, machine learning and survival analysis.

JLDR, ASD, JG, RI as specialized dermatologists have diagnosed and doing the follow up of all the patients included in this study. Moreover, they have analyzed and interpreted the data according the clinical significance of the results.

AAp and GPY analyzed and interpreted all the clinical and experimental data and they have contributed in writing the manuscript.

PAE and CP: Samples curation and revision of clinical data and tumor evolution of patients.

All the authors read and approved the final manuscript.

## Notes

FM, SL and JR contributed equally to this work

## Consent for publication

Manuscript is approved by all authors for publication.

## Figure Legends

**Figure S1.**
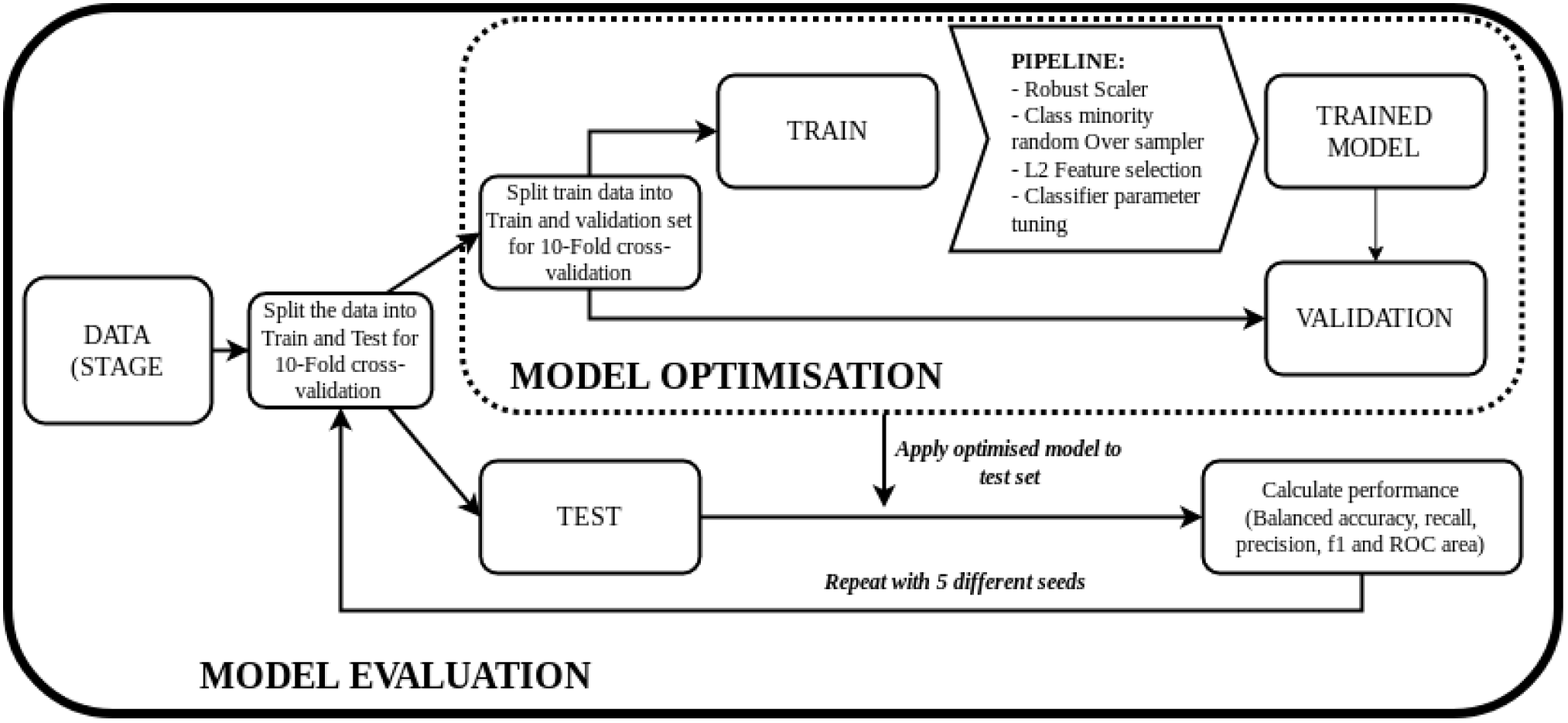
Workflow of the machine learning analysis.

**Figure S2.**
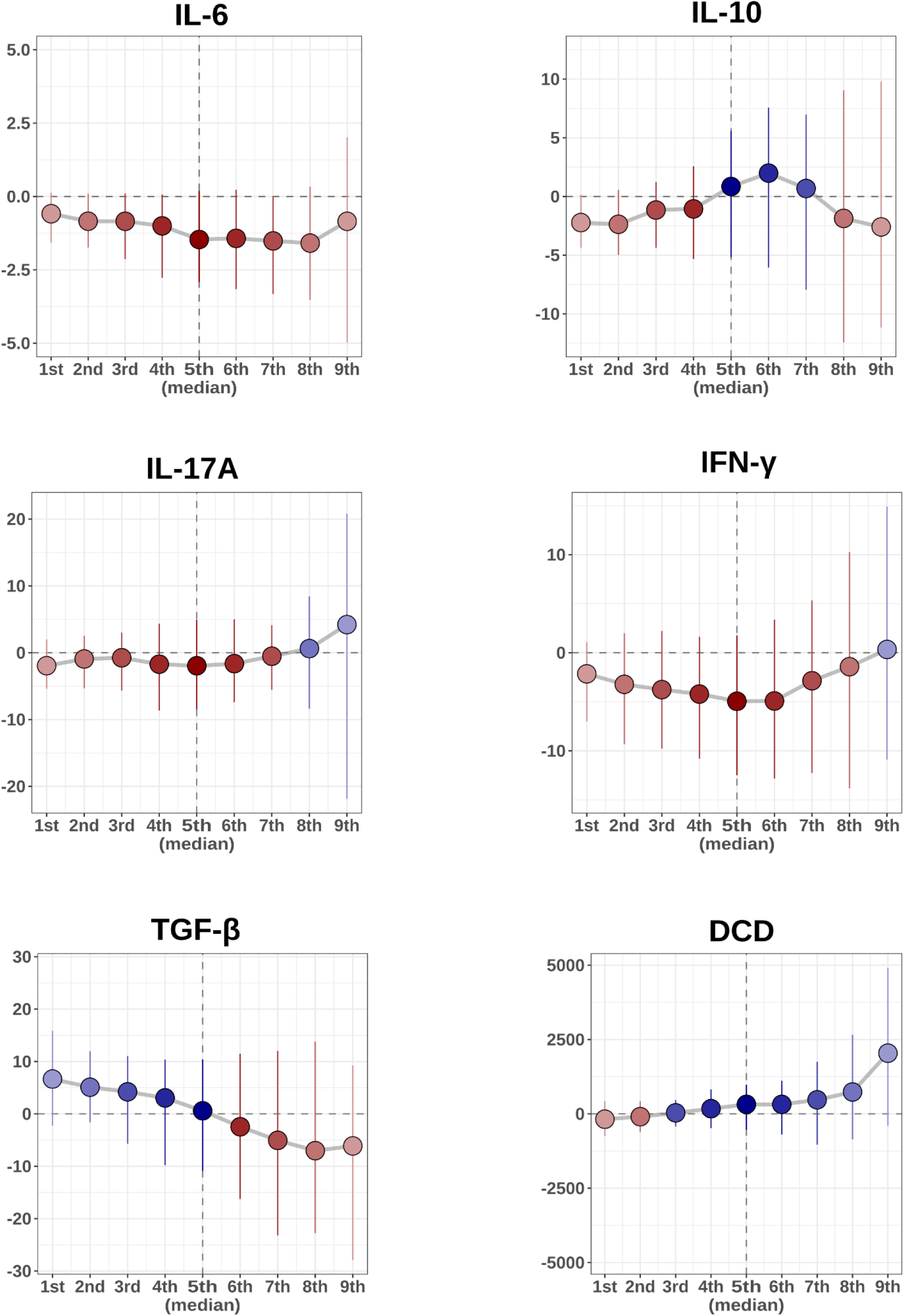
Shift function for the rest of variables of interest, displaying the difference between the deciles of the subgroup of disease-free and metastatic subjects. Positive values of the shift function are in blue, corresponding to larger decile values in the disease-free group than in the metastatic group, while red values illustrate the opposite scenario.

**Figure S3.**
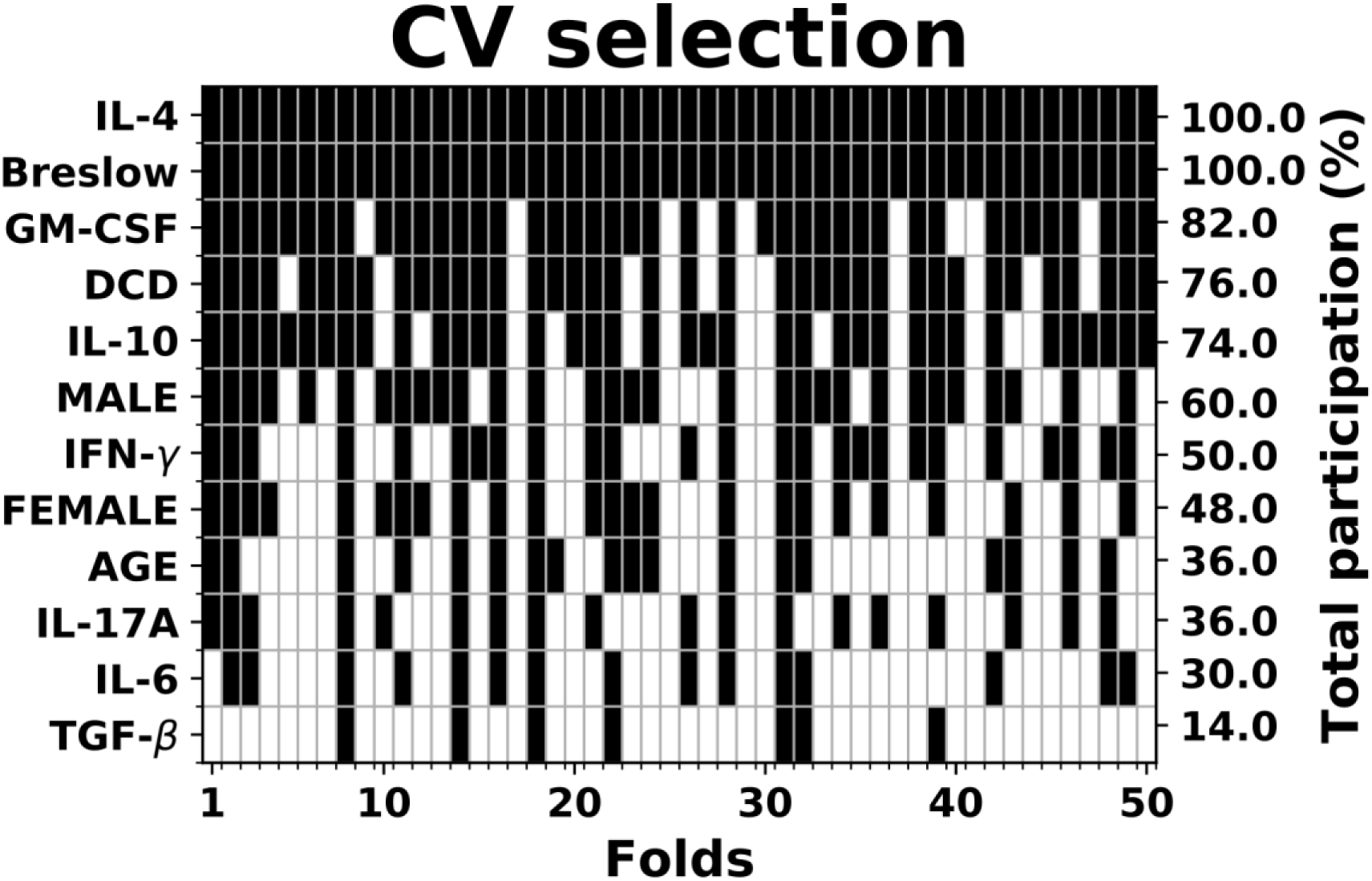
Participation across the folds provided by the feature selection step in the inner cross-validation loop when confounding variables age and sex (separated in male and female categories) are incorporated. Black colors in each column denote the predictors that were included in the final logistic regression model in each of these folds. Data from the early-stage melanoma cohort (n=323).

**Table S1.** Train and test prediction scores in the three domains of variables for the battery of algorithms used.

| **BRESLOW THICKNESS** |  |  |  |  |  |  |  |  |
| --- | --- | --- | --- | --- | --- | --- | --- | --- |
|  | **Train**<br>**Balanced**<br>**accuracy** | **Test**<br>**Balanced**<br>**accuracy** | **Train**<br>**Recall** | **Test**<br>**Recall** | **Train**<br>**Precision** | **Test**<br>**Precision** | **Train**<br>**Area**<br>**ROC** | **Test**<br>**Area**<br>**ROC** |
| **LR** | 74.92 | 73.02 | 72.65 | 69.50 | 52.98 | 51.69 | 83.45 | 83.72 |
| **SVM** | 74.03 | 73.57 | 65.65 | 64.94 | 56.37 | 58.4 | 82.28 | 81.51 |
| **DT** | 77.93 | 72.56 | 76.81 | 69.06 | 57.07 | 53.5 | 84.07 | 78.72 |
| **NB** | 72.66 | 72.56 | 54.76 | 54.67 | 66.65 | 69.17 | 83.08 | 83.28 |
| **KNN** | 78.69 | 73.26 | 80.32 | 72.94 | 54.99 | 49.67 | 87.65 | 79.17 |
| **CYTOKINES + DCD** |  |  |  |  |  |  |  |  |
|  | **Train**<br>**Balanced**<br>**accuracy** | **Test**<br>**Balanced**<br>**accuracy** | **Train**<br>**Recall** | **Test**<br>**Recall** | **Train**<br>**Precision** | **Test**<br>**Precision** | **Train**<br>**Area**<br>**ROC** | **Test**<br>**Area**<br>**ROC** |
| **LR** | 65.75 | 59.98 | 62.69 | 52.73 | 35.03 | 29.83 | 71.98 | 65.42 |
| **SVM** | 69.04 | 58.19 | 53.78 | 35.04 | 47.96 | 33.28 | 75.25 | 62.98 |
| **DT** | 73.57 | 57.79 | 70.88 | 47.73 | 46.58 | 30.12 | 78.74 | 58.38 |
| **NB** | 60.62 | 55.52 | 43.01 | 35.40 | 45.60 | 31.44 | 65.9 | 58.50 |
| **KNN** | 72.21 | 56.78 | 74.96 | 52.07 | 39.68 | 26.80 | 79.69 | 56.16 |
| **BRESLOW THICKNESS + CYTOKINES + DCD** |  |  |  |  |  |  |  |  |
|  | **Train**<br>**Balanced**<br>**accuracy** | **Test**<br>**Balanced**<br>**accuracy** | **Train**<br>**Recall** | **Test**<br>**Recall** | **Train**<br>**Precision** | **Test**<br>**Precision** | **Train**<br>**Area**<br>**ROC** | **Test**<br>**Area**<br>**ROC** |
| **LR** | 82.38 | 80.37 | 79.41 | 77.53 | 59.40 | 57.38 | 90.95 | 89.22 |
| **SVM** | 80.15 | 77.56 | 73.18 | 68.60 | 60.44 | 60.02 | 87.94 | 85.99 |
| **DT** | 86.54 | 76.03 | 87.90 | 70.8 | 62.66 | 53.87 | 92.68 | 85.46 |
| **NB** | 79.73 | 77.79 | 69.09 | 66.02 | 65.68 | 63.46 | 88.45 | 85.62 |
| **KNN** | 85.02 | 74.52 | 86.26 | 70.73 | 58.88 | 46.97 | 92.66 | 81.99 |

## References

1. Tracey EH, Vij A. Updates in Melanoma. Dermatol Clin 2019; 37(1):73–82.

2. Whiteman DC, Green AC, Olsen CM. The Growing Burden of Invasive Melanoma: Projections of Incidence Rates and Numbers of New Cases in Six Susceptible Populations through 2031. J Invest Dermatol 2016; 136(6):1161–1171.

3. Kashani-Sabet M, Nosrati M, Miller JR 3rd, Sagebiel RW, Leong SPL, Lesniak A, Tong S, Lee SJ, Kirkwood JM. Prospective Validation of Molecular Prognostic Markers in Cutaneous Melanoma: A Correlative Analysis of E1690. Clin Cancer Res 2017; 23(22):6888–6892.

4. Kashani-Sabet M. Molecular markers in melanoma. Br J Dermatol 2014; 170(1):31–5.

5. Foth M, Wouters J, de Chaumont C, Dynoodt P, Gallagher WM. Prognostic and predictive biomarkers in melanoma: an update. Expert Rev Mol Diagn 2016; 16(2):223–37.

6. Stiegel E, Xiong D, Ya J, Funchain P, Isakov R, Gastman B, Vij A. Prognostic value of sentinel lymph node biopsy according to Breslow thickness for cutaneous melanoma. J Am Acad Dermatol. 2018; 78(5):942–948.

7. Balch CM, Gershenwald JE, Soong SJ, Thompson JF, Atkins MB, Byrd DR, Buzaid AC, Cochran AJ, Coit DG, Ding S, Eggermont AM, Flaherty KT, Gimotty PA, Kirkwood JM, McMasters KM, Mihm MC Jr, Morton DL, Ross MI, Sober AJ, Sondak VK. Final version of 2009 AJCC melanoma staging and classification. J Clin Oncol 2009; 27(36):6199–206.

8. Elmore JG, Elder DE, Barnhill RL, Knezevich SR, Longton GM, Titus LJ, Weinstock MA, Pepe MS, Nelson HD, Reisch LM, Radick AC, Piepkorn MW. Concordance and Reproducibility of Melanoma Staging According to the 7th vs 8th Edition of the AJCC Cancer Staging Manual. JAMA Netw Open 2018; 1(1).

9. Weiss SA, Hanniford D, Hernando E, Osman I. Revisiting determinants of prognosis in cutaneous melanoma. Cancer 2015; 121(23):4108–23.

10. Gogas H, Eggermont AMM, Hauschild A, Hersey P, Mohr P, Schadendorf D, Spatz A, Dummer R. Biomarkers in melanoma. Ann Oncol 2009; 20 Suppl 6:vi8–13.

11. Palmer SR, Erickson LA, Ichetovkin I, Knauer DJ, Markovic SN. Circulating serologic and molecular biomarkers in malignant melanoma. Mayo Clin Proc 2011; 86(10):981–90.

12. Johdi NA, Mazlan L, Sagap I, Jamal R. Profiling of cytokines, chemokines and other soluble proteins as a potential biomarker in colorectal cancer and polyps. Cytokine 2017; 99:35–42.

13. Obraztsov IV, Shirokikh KE, Obraztsova OI, Shapina MV, Wang MH, Khalif IL. Multiple Cytokine Profiling: A New Model to Predict Response to Tumor Necrosis Factor Antagonists in Ulcerative Colitis Patients. Inflamm Bowel Dis 2019; 25(3):524–531.

14. D’Angelo C, Reale M, Costantini E, Di Nicola M, Porfilio I, de Andrés C, Fernández-Paredes L, Sánchez-Ramón S, Pasquali L. Profiling of Canonical and Non-Traditional Cytokine Levels in Interferon-β-Treated Relapsing-Remitting-Multiple Sclerosis Patients. Front Immunol 2018; 9:1240.

15. Nevala WK, Vachon CM, Leontovich AA, Scott CG, Thompson MA, Markovic SN; Melanoma Study Group of the Mayo Clinic Cancer Center. Evidence of systemic Th2-driven chronic inflammation in patients with metastatic melanoma. Clin Cancer Res 2009; 15(6):1931–9.

16. Boyano MD, Garcia-Vázquez MD, López-Michelena T, Gardeazabal J, Bilbao J, Cañavate ML, García de Galdeano A, Izu R, Díaz-Ramón L, Raton JA, Díaz-Pérez JL. Soluble interleukin-2 receptor, intercellular adhesion molecule-1 and interleukin-10 serum levels in patients with melanoma. Br J Cancer 2000; 83(7):847–52.

17. Boyano MD, García-Vázquez MD, Gardeazabal J, García de Galdeano A, Smith-Zubiaga I, Cañavate ML, Raton JA, Bilbao I, Díaz-Pérez JL. Serum-soluble IL-2 receptor and IL-6 levels in patients with melanoma. Oncology 1997; 54(5):400–6.

18. Ma YF, Chen C, Li D, Liu M, Lv ZW, Ji Y, Xu J. Targeting of interleukin (IL)-17A inhibits PDL1 expression in tumor cells and induces anticancer immunity in an estrogen receptor-negative murine model of breast cancer. Oncotarget 2017; 8(5):7614–7624.

19. Bhattacharya P, Thiruppathi M, Elshabrawy HA, Alharshawi K, Kumar P, Prabhakar BS. GM-CSF: An immune modulatory cytokine that can suppress autoimmunity. Cytokine 2015; 75(2):261–71.

20. Singel KL, Segal BH. Neutrophils in the tumor microenvironment: trying to heal the wound that cannot heal. Immunol Rev 2016; 273(1):329–43.

21. Reggiani F, Labanca V, Mancuso P, Rabascio C, Talarico G, Orecchioni S, Manconi A, Bertolini F. Adipose Progenitor Cell Secretion of GM-CSF and MMP9 Promotes a Stromal and Immunological Microenvironment That Supports Breast Cancer Progression. Cancer Res 2017; 77(18):5169–5182.

22. Wang TT, Zhao YL, Peng LS, Chen N, Chen W, Lv YP, Mao FY, Zhang JY, Cheng P, Teng YS, Fu XL, Yu PW, Guo G, Luo P, Zhuang Y, Zou QM. Tumour-activated neutrophils in gastric cancer foster immune suppression and disease progression through GM-CSF-PD-L1 pathway. Gut 2017; 66(11):1900–1911.

23. Ortega-Martínez I, Gardeazabal J, Erramuzpe A, Sanchez-Diez A, Cortés J, García-Vázquez MD, Pérez-Yarza G, Izu R, Luís Díaz-Ramón J, de la Fuente IM, Asumendi A, Boyano MD. Vitronectin and dermcidin serum levels predict the metastatic progression of AJCC I-II early-stage melanoma. Int J Cancer 2016; 139(7):1598–607.

24. Zeth K, Sancho-Vaello E. The Human Antimicrobial Peptides Dermcidin and LL-37 Show Novel Distinct Pathways in Membrane Interactions. Front Chem 2017; 5: 86.

25. Paulmann M, Arnold T, Linke D, Özdirekcan S, Kopp A, Gutsmann T, Kalbacher H, Wanke I, Schuenemann VJ, Habeck M, Bürck J, Ulrich AS, Schittek B. Structure-Activity Analysis of the Dermcidin-derived Peptide DCD-1L, an Anionic Antimicrobial Peptide Present in Human Sweat. J Biol Chem 2012; 287(11): 8434–43.

26. Wagenmakers, E-J. A practical solution to the pervasive problems of p values. Psychon. Bull. Rev 2007; 14: 779–804.

27. Mohammadpour A, Derakhshan M, Darabi H, Hedayat P, Momeni M. Melanoma: Where we are and where we go. J Cell Physiol 2019; 234(4):3307–3320.

28. Svedman FC, Pillas D, Taylor A, Kaur M, Linder R, Hansson J. Stage-specific survival and recurrence in patients with cutaneous malignant melanoma in Europe - a systematic review of the literature. Clin Epidemiol 2016; 8:109–22.

29. Rutkowski P, Lugowska I. Follow-up in melanoma patients. Memo 2014; 7(2):83–86.

30. Karagiannis P, Fittall M, Karagiannis SN. Evaluating biomarkers in melanoma. Front Oncol 2015; 4:383.

31. Qiu F, Qiu F, Liu L, Liu J, Xu J, Huang X. The Role of Dermcidin in the Diagnosis and Staging of Hepatocellular Carcinoma. Genet Test Mol Biomarkers 2018; 22(4):218–223.

32. Trzoss L, Fukuda T, Costa-Lotufo LV, Jimenez P, La Clair JJ, Fenical W. Seriniquinone, a selective anticancer agent, induces cell death by autophagocytosis, targeting the cáncer-protective protein dermcidin. Proc Natl Acad Sci U S A 2014; 111(41):14687–92.

33. Wang M, Zhao J, Zhang L, Wei F, Lian Y, Wu Y, Gong Z, Zhang S, Zhou J, Cao K, Li X, Xiong W, Li G, Zeng Z, Guo C. Role of tumor microenvironment in tumorigenesis. J Cancer. 2017; 8(5):761–773.

34. Terren I, Orrantia A, Vitalle J, Zenarruzabeitia O and Borrego F. NK cell metabolism and tumor microenvironment. Front Immunology 2019; 10, 2278.

35. Jiang X, Wang J, Deng X, Xiong F, Ge J, Xiang B, Wu X, Ma J, Zhou M, Li X, Li Y, Li G, Xiong W, Guo C, Zeng Z. Role of the tumor microenvironment in PD-L1/PD-1-mediated tumor immune escape. Mol Cancer 2019; 18(1):10.

36. Ito SE, Shirota H, Kasahara Y, Saijo K, Ishioka C. IL-4 blockade alters the tumor microenvironment and augments the response to cancer immunotherapy in a mouse model. Cancer Immunol Immunother 2017; 66(11):1485–1496.

37. Setrerrahmane S and Xu H. Tumor-related interleukins: old validated targets for new anti-cancer drug development. Mol Cancer 2017; 16(1):153.

38. Surcel M, Constantin C, Caruntu C, Zurac S, Neagu M. Inflammatory Cytokine Pattern Is Sex-Dependent in Mouse Cutaneous Melanoma Experimental Model. J Immunol Res 2017; 2017:9212134.

39. Jobe NP, Živicová V, Mifková A, Rösel D, Dvořánková B, Kodet O, Strnad H, Kolář M, Šedo A, Smetana K Jr, Strnadová K, Brábek J, Lacina L. Fibroblasts potentiate melanoma cells in vitro invasiveness induced by UV-irradiated keratinocytes. Histochem Cell Biol 2018; 149(5):503–516.

40. Yu SH, Maynard JP, Vaghasia AM, De Marzo AM, Drake CG, Sfanos KS. A role for paracrine interleukin-6 signaling in the tumor microenvironment in prostate tumor growth. Prostate 2019; 79(2):215–222.

41. Gocheva V, Wang HW, Gadea BB, Shree T, Hunter KE, Garfall AL, Berman T, Joyce JA. IL-4 induces cathepsin protease activity in tumor-associated macrophages to promote cancer growth and invasion. Genes Dev 2010; 24(3):241–55.

42. Hong IS. Stimulatory versus suppressive effects of GM-CSF on tumor progression in multiple cancer types. Exp Mol Med 2016; 48(7):e242.

43. Cliff, N. Dominance statistics: Ordinal analyses to answer ordinal questions. Psychol Bull 1993; 114(3):494–509.

44. Benjamini Y, Hochberg Y. Controlling the False Discovery Rate: A Practical and Powerful Approach to Multiple Testing. J. Royal Stat. Soc, 1995; 57(1):289–300.

45. Rand R. Wilcox. Introduction to Robust Estimation and Hypothesis Testing, 3rd ed., California, Elsevier, 2012. 608p

46. Rousselet GA, Pernet CR, Wilcox RR. Beyond differences in means: robust graphical methods to compare two groups in neuroscience. Eur J Neurosci 2017; 46(2):1738–1748.

47. Pedregosa F, Varoquaux G, Gramfort A, Michel V, Thirion B, Grisel O, Blondel M, Prettenhofer P, Weiss R, Dubourg V, Vanderplas J, Passos A, et al. Scikit-learn: Machine Learning in Python. J Mach Learn Res 2011; 12:2825–30.

48. Lemaître G, Nogueira F, Aridas CK. Imbalanced-learn: A Python Toolbox to Tackle the Curse of Imbalanced Datasets in Machine Learning. J Mach Learn Res 2017; 18:1–5.

49. Ilker Unal. Defining an Optimal Cut-Point Value in ROC Analysis: An Alternative Approach. Comput Math Methods Med 2017; 2017:14.

50. Davidson-Pilon C, Kalderstam J, Kuhn B, Fiore-Gartland A, Moneda L, Zivich P, Parij A, Stark K, Anton S, Besson, Jona, Harsh Gadgil, et al. Software Open Access CamDavidsonPilon/lifelines: v0.14.3. 2018.

